# CRISPR/Cas9-based genome editing in the silverleaf whitefly (*Bemisia tabaci*)

**DOI:** 10.1101/2020.03.18.996801

**Authors:** Chan C. Heu, Francine M. McCullough, Junbo Luan, Jason L. Rasgon

## Abstract

*Bemisia tabaci* cryptic species Middle East-Asia Minor I (MEAM1) is a serious agricultural polyphagous insect pest, and vector of numerous plant viruses, causing major worldwide economic losses. *B. tabaci* control is limited by lack of robust gene editing tools. Gene editing is difficult in *B. tabaci* due to small embryos that are technically challenging to inject, and which have high mortality post-injection. We developed a CRISPR/Cas9 gene editing protocol based on injection of vitellogenic adult females rather than embryos (“ReMOT Control”). We identified an ovary-targeting peptide ligand (“BtKV”) that, when fused to Cas9 and injected into adult females, transduced the ribonucleoprotein complex to the germline, resulting in efficient, heritable editing of the offspring genome. In contrast to embryo injection, adult injection is easy and does not require specialized equipment. Development of easy-to-use gene editing protocols for *B. tabaci* will allow researchers to apply the power of reverse genetic approaches to this species and will lead to novel control methods for this devastating pest insect.

## Introduction

*Bemisia tabaci* cryptic species Middle East-Asia Minor I (MEAM1) is a widely distributed invasive agricultural pest that causes the economic loss of billions of dollars in crop damages worldwide (1, 2). *B. tabaci* is polyphagous with a broad host range. This insect feeds on plant phloem sap through its life cycle using piercing-sucking mouthparts and can cause direct damage to the plants. The honeydew excretions from whiteflies promote fungus growth that can reduce photosynthesis and crop yields. Finally, *B. tabaci* is an important vector of numerous plant viruses that affect economically critical crop species. *B. tabaci* is the only known vector of begomoviruses, a family of plant viruses known to cause plant diseases and adversely affect crop yield (2). Current control methods for *B. tabaci* are insecticides, and limited use of biological control (3, 4). Plant-mediated RNAi targeting begomoviruses or the whitefly vector has shown some promise in laboratory settings but has not been translated to field applications (5 - 8). Although RNAi can be effective at reducing gene transcription, efficacy can be highly variable depending on the gene and tissue of interest. Thus far, the lack of tools to genetically manipulate whitefly hinders the screening of potential genetic targets that can be used to design agricultural control strategies.

The economic importance of *B. tabaci* demands new methods to control this devastating pest insect. Gene editing, using CRISPR/Cas9, has formed the foundation for novel control strategies for insect vectors of human diseases (9 - 13) and plant diseases (14), but lack of gene editing techniques for *B. tabaci* is a significant barrier to the application of gene editing for basic biological studies and control of this insect. Arthropod gene editing by CRISPR/Cas9 is usually performed by injecting gene-editing materials into pre-blastoderm embryos, but the exceedingly small size of *B. tabaci* embryos (0.2mm) and high mortality of injected eggs makes this technically challenging. A method has recently been developed called “<u>Re</u>MOT Control” (<u>Re</u>ceptor-<u>M</u>ediated <u>O</u>vary <u>T</u>ransduction of <u>C</u>argo) which circumvents the need to inject embryos, but rather uses a small ovary-targeting peptide to directly transduce the Cas9 ribonucleoprotein complex (RNP; the Cas9 protein complexed with a guide RNA) into the developing ovaries upon injection into the hemolymph of adult female insects. ReMOT Control has been shown to efficiently edit the germline of the mosquitoes *Aedes aegypti* and *Anopheles stephensi* (9, 10).

Here, we develop an efficient ReMOT Control CRISPR/Cas9-based adult injection protocol for gene editing in *B. tabaci*. The development of robust gene editing methodologies for this species opens the power of CRISPR and reverse genetic approaches to study the biology and develop new control strategies for this important economic pest.

## Methods and Materials

### Bemisia tabaci colony maintenance

A colony of *B. tabaci* MEAM1 (originally collected from a poinsettia plant in Ithaca, NY) was maintained in a growth chamber set at 28**°**C +/-2**°**C, 14L/10D photoperiod and 50% relative humidity in 24 × 24 × 24 inch cages with organza access sleeves. Insects were maintained on organic soybean (var. Viking, Johnny’s Selected Seeds), organic radish (var. *Cherry* Belle, Burpee Seeds and Plants), and/or gerbera daisy (var. *Garvinea*, Burpee Seeds and Plants). Plants were grown in a separate growth chamber using the same parameters as the insect growth chamber. Plants were grown in 6-inch pots using enriched potting media, with liquid fertilizer (Peter’s Excel Cal-Mag 15-5-15) applied twice a month.

### Plasmid construction and protein expression

Based on a predicted 24 aa vitellogenin binding sequence from the crustacean *Macro-brachiumrosen bergii* (15) we identified the homologous sequence from multiple *B. tabaci* vitellogenin genes (XP_018897090, XP_018912902, XP_018897089), and from the mosquitoes *Ae. aegypti* (AAA994861.1), and *An. gambiae* (AAF82131.1). We identified a lysine conserved across *B. tabaci* sequences and a fully conserved valine residue that defined the ends of the targeting sequence (“BtKV”) (Supplementary Figure 1). pET28A-BtKV-Cas9 was constructed from pET28-P2C-Cas9 (10) and a gblock (IDT) of BtKV-mCherry. pET28A-P2C-Cas9 was digested with restriction enzymes BamHI and SalI. A gblock of BtKV fused to mCherry was synthesized and inserted into the digested pET28-P2C-Cas9 using In-Fusion Cloning (Takara). Sanger sequencing was used to verify the sequence. pET28A-BtKV-mCherry-Cas9 was further digested with either SalI or XhoI to remove mCherry or Cas9 respectively, gel purified, and re-ligated using T4 ligase to obtain the plasmids pET28A-BtKV-Cas9 or pET28A-BtKV-mCh. Plasmids were transformed into BL21(DE3) cells for BtKV-Cas9/mCh expression following standard protein expression protocols. P2C-Cas9 and Cas9 were expressed using plasmids as previously described (10). Briefly, a 5 mL starter culture grown overnight at 37°C with shaking at 225 rpm was used to inoculate 50 mL Terrific Broth (Invitrogen) at a 1:100 dilution. When the culture reached an OD_600_ of 0.4-0.8, it was induced with isopropyl-β-D-1-thiogalactopyranoside (IPTG) at [500 μM] concentration at 20°C overnight. Cultures were pelleted, resuspended in lysis buffer (20 mM Tris-HCl pH 8.0, 300 mM NaCl, 20 mM imidazole) and sonicated. Cell debris was pelleted and the supernatant collected and used for immobilized metal affinity chromatography using Ni-NTA agarose beads (Qiagen) and agitated for at least 2 hours at 4°C. Subsequently, the beads were separated in a chromatography column and washed with lysis buffer allowing gravity flow. Proteins were eluted (20 mM Tris-HCl pH 8.0, 300 mM NaCl, 250 mM imidazole) and dialyzed in 20 mM Tris-HCl pH 8.0, 300 mM KCl, 500 μM phenylmethylsulfonyl fluoride. Proteins were concentrated using an Amicon Ultra-0.5 mL centrifugal filter device with a cutoff of 100 kDa (Millipore Sigma).

### Generating guide RNAs

sgRNA were generated following the protocols found in (11). The 20 nucleotide guide sequences were designed against exon 2, exon 3, and exon 5 of the *B. tabaci white* gene (NW_017550151, region 408016-472564) both manually and by CRISPOR (16). Primers and guide RNA sequences are listed in Supplementary Table 1.

### Injection protocol for Bemisia tabaci

For initial experiments *B. tabaci* adults of unknown age and mating status were aspirated from the colony cage into 2-dram screw cap vials, placed in ice to anesthetize the insects, sexed, and transferred to a standard plain glass slide with double-sided tape on a chill table. For later experiments we established a synchronous colony so that we could control for the age of injected insects (see below). For localization of BtKV fusion protein into the embryos, female *B. tabaci* were injected in the abdomen with BtKV-mCherry fusion protein (3 mg/ml) or PBS control using quartz capillary needles. Samples were collected 24 hours post injection and ovaries dissected onto a concavity microscope slide with a glass coverslip and imaged by epifluorescence microscopy.

To generate mutants, the injection mixture consisted of the RNP complex of BtKV-Cas9 (3 mg/mL) with a mix of all sgRNAs (3 mg/mL) at a 1:3 molar ratio, and 1/3^rd^ the volume of saponin [4, 8, or 16 μg/mL] as an endosome escape reagent (or buffer as a control) (9, 10). Five sgRNA multiplexed were used at a concentration of 250 ng/μL each and 1.25 μg/μL total. Two groups of females were collected: 1) Females of unknown age and unknown mated status were collected at random and 2) females less than 24 hours post emergence (hpe) were collected to reduce their chances of having already developed eggs (18). The females were injected and placed on a piece of soybean leaflet in a petri dish with a moist paper towel wrapped around the stem and another paper towel on the bottom plate to prolong the life of the leaflet. Water and liquid fertilizer were added as needed. Females were removed from the leaflet 2 weeks post injection. The petri dishes were incubated at 28°C, 16L/8D cycle, in a humidified Heratherm IMH750-S reach-in chamber, and the progeny of injected females screened visually for altered eye color in nymphs or adults using an Olympus SZX7 stereomicroscope. Mutations in insects showing altered eye color were confirmed by PCR using PHIRE Tissue Direct PCR Master Mix (Thermo Fischer Scientific), the amplicon was cloned into pJET1.2, and clones were submitted for Sanger sequencing.

### Heritability crosses

Crosses of mutant males and wild-type females were performed (Fig. 4) to demonstrate heritability of the mutant allele and phenotype. In brief, G0 mutant males resulting from injection of G-1 females were interbred with wild-type G0 females. Resulting G1’s were sexed and separated approximately 24 hpe. G1 females were backcrossed to a G0 mutant male to generate G2 offspring. Insects were screened for eye color in the nymphal stage.

**Figure 1.**
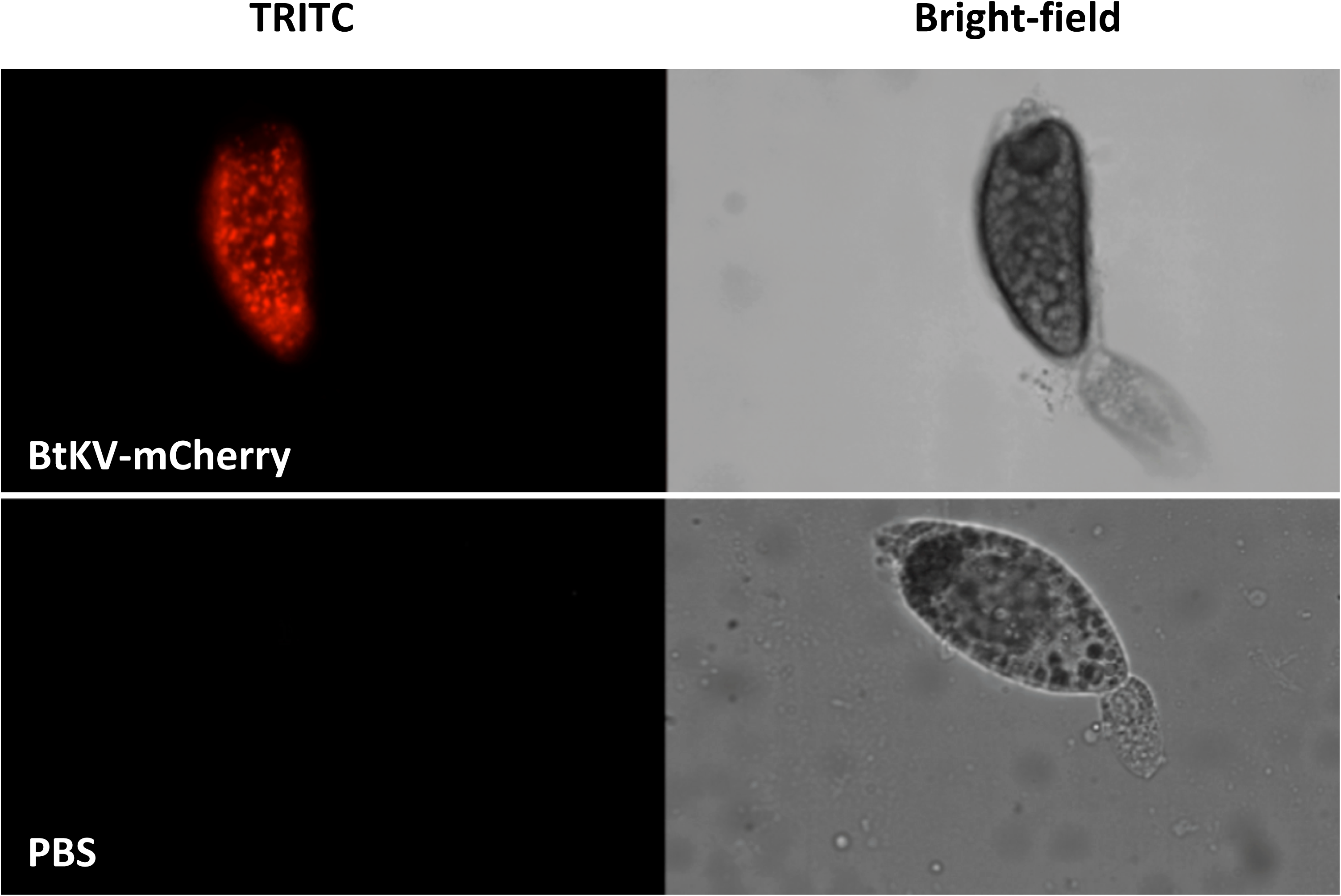
Targeting of molecular cargo to *B. tabaci* ovaries. The BtKV peptide targets molecular cargo (mCherry-Cas9 fusion protein) into *B. tabaci* ovaries and is visible in dissected mature oocytes 24 hours post-injection (top). No signal is visible in control ovaries (bottom).

**Figure 2.**
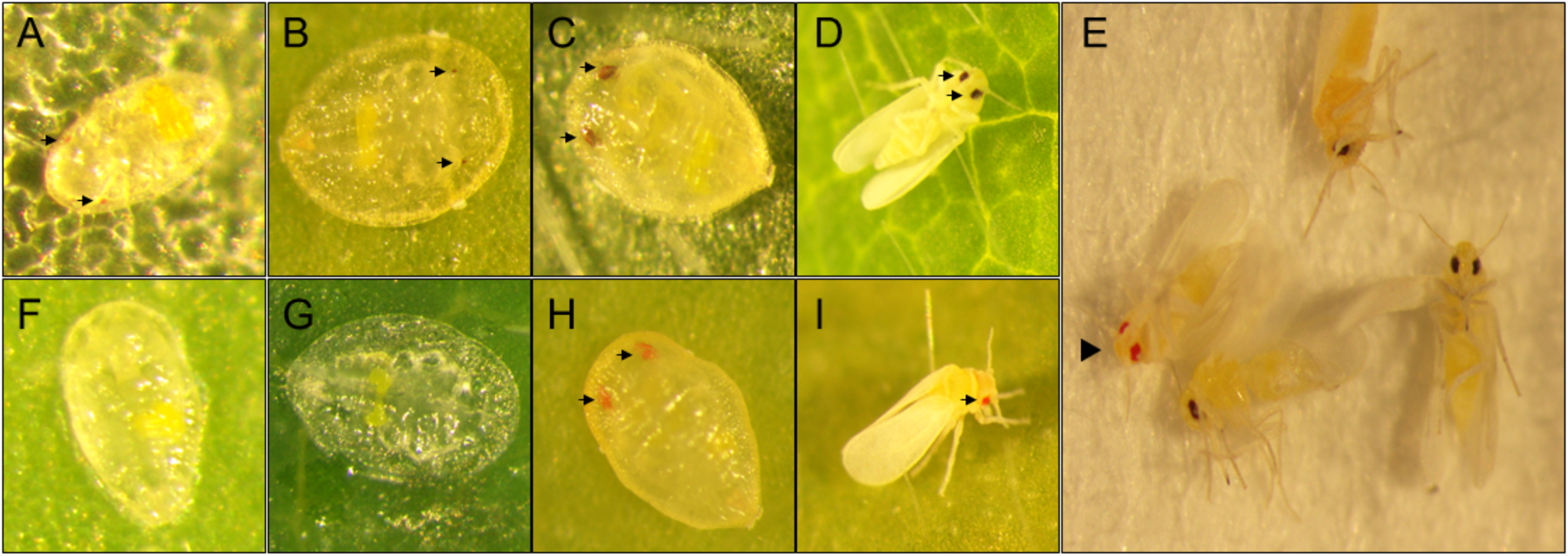
Wild-type and mutant *B. tabaci*. Panels A, B, C, and D show wild-type from 2^nd^, 3^rd^, and 4^th^ instar nymphs, and adult, respectively. Panels F, G, H, and I show matched-stage mutant individuals resulting from ReMOT Control editing of the *B. tabaci white* gene. Black arrows point to eye(s) of wild-type and mutants. Panel E shows both mutant and wild-type in the same image. Black arrowhead points to mutant adult.

**Figure 3.**
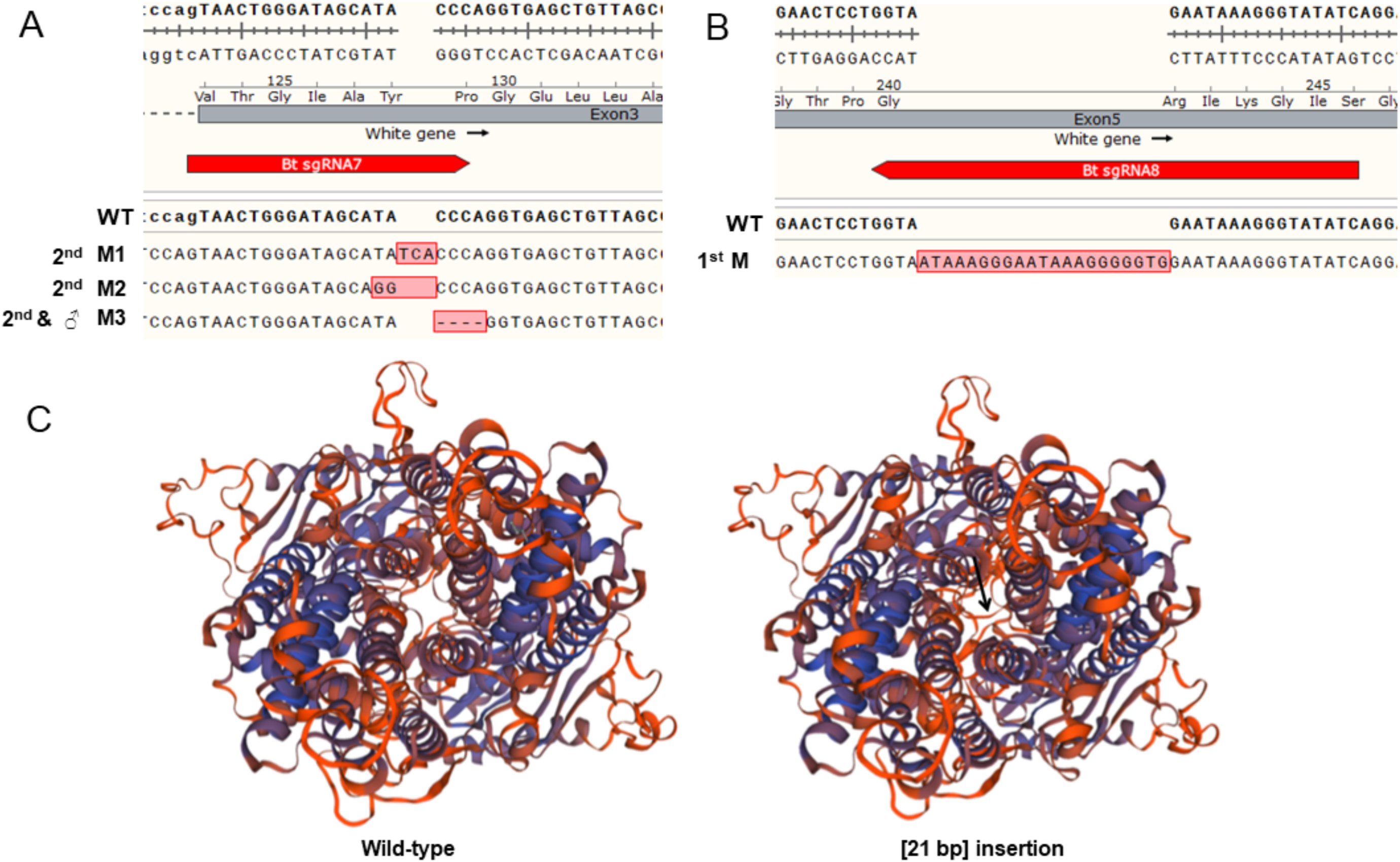
Mutant alleles and homology modeling of the [21bp] insertion. A) The mutant alleles (M1; 3 nt insertion and M2; 2 nt substitution, M3; 4 nt deletion) identified at exon 3 were compared to the wild-type sequence. 2^nd^ and ♂ indicates the instar/life stage the mutation was identified in (2 adult males shared this mutation). B) The 21bp insertion (M) at exon 5 in a 1^st^ nymphal instar. C) Predicted 3D structure of the wild-type *B. tabaci white* protein (left) compared to the structure of the in-frame 21bp insertion mutant (right). The insertion is predicted to cause a loop (arrow) across the channel pore, sterically hindering function of the transporter, leading to a null mutation.

**Figure 4.**
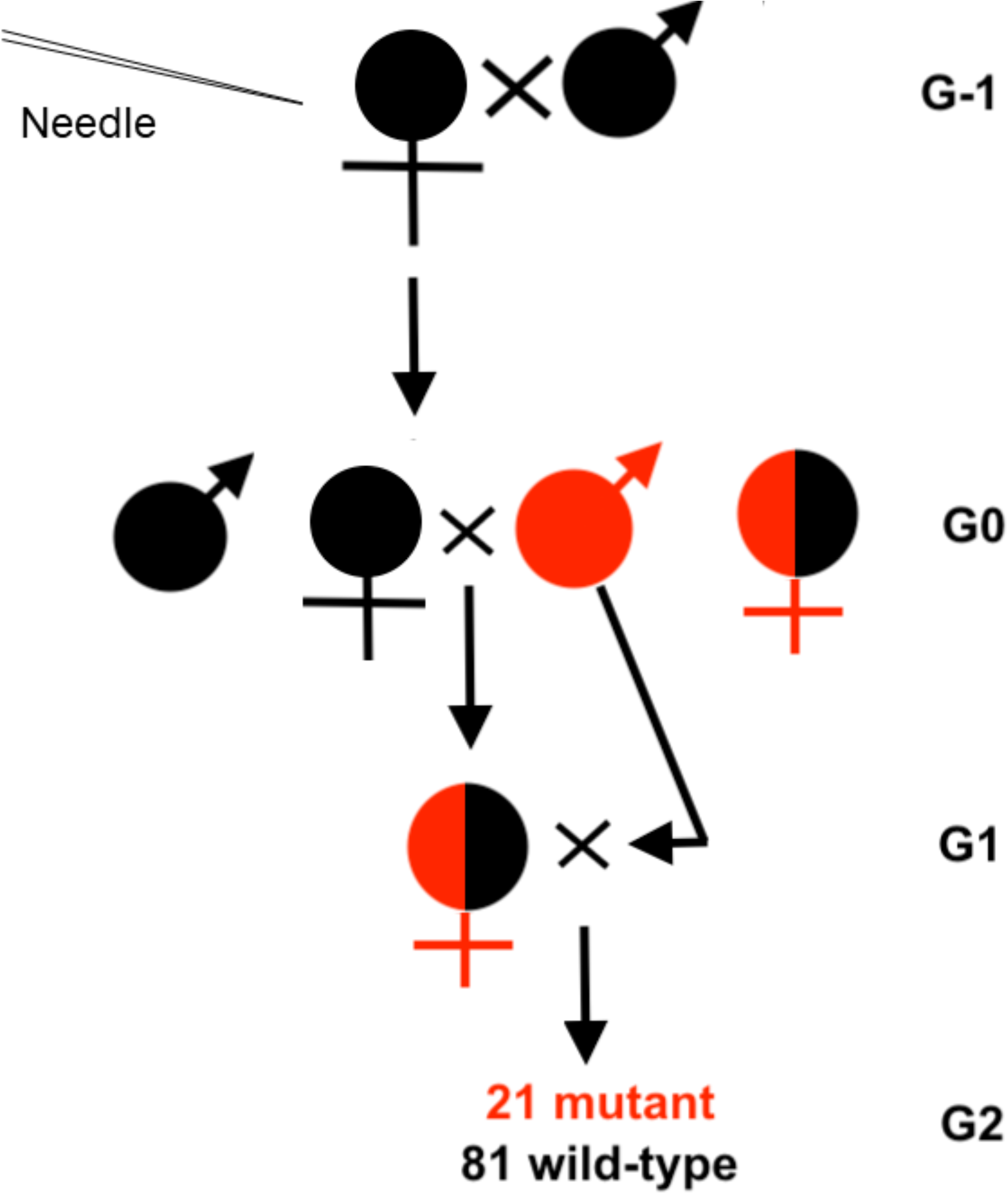
Crossing scheme to identify heritability. Injected females (G-1) can produce wild-type males and females (black), mutant males (red), and/or hemizygous females (red/black) in G0. The resulting mutant males were crossed with the female siblings to generate the G1. The G1 hemizygous females were backcrossed with G0 mutant males to generate G2 offspring which were scored as juveniles for white or wild-type eyes.

### Homology modeling of mutated proteins

A homology model of one in-frame insertion mutant [21bp] was generated using the Swiss Model homology server (swissmodel.expasy.org). The structure of the *B. tabaci white* gene has not been empirically solved, so a model of the wild-type protein was first generated against the database using the server, the result of which was used as template to model the [21bp] insertion mutant.

## Results

### *Identification and validation of* thse BtKV *ovary targeting ligand*

While the P2C ligand works to target cargo to the ovaries of multiple mosquito species (9, 10), it did not function robustly in *B. tabaci* (Table 1). We therefore designed a new targeting peptide suitable for this insect species based on its endogenous vitellogenin protein (“BtKV”; KPYGVYKTMEDSV) (see methods). *B. tabaci* oocyte development is asynchronous with stage I, II, III, and IV at 11 days post eclosion or older (19) where oocytes are always developing. Developmental phase II oocytes have the highest amount of vitellogenin uptake via endocytosis and nutrient cord (19). After injection of BtKV-mCherry into reproductive adult *B. tabaci* females, ovaries were dissected after 24 hours and visualized for red fluorescence. We observed red fluorescence in the developing oocytes from females injected with BtKV-mCherry fusion protein but not PBS-injected controls (Fig. 1).

**Table 1.** CRISPR editing efficiency in *B. tabaci*.

| Injection | Saponin<br>($\mu\text{g/mL}$) | N females<br>injected | Sampling<br>method | N Offspring<br>screened | N Offspring<br>edited | KO/injected<br>(%) | KO/offspring<br>(%) | KO/offspring (%)<br>Sex Ratio<br>Corrected |
| --- | --- | --- | --- | --- | --- | --- | --- | --- |
| Cas9 | 0 | 31 | Random | 70 | 0 | 0 | 0 | 0 |
| Cas9 | 4 | 28 | Random | 218 | 0 | 0 | 0 | 0 |
| P2C-Cas9 | 0 | 20 | Random | 54 | 0 | 0 | 0 | 0 |
| P2C-Cas9 | 4 | 13 | Random | 74 | 0 | 0 | 0 | 0 |
| BtkV-Cas9 | 0 | 42 | Random | 961 | 1 | 2.38% | 0.10% | 0.39% |
| BtkV-Cas9 | 4 | 47 | Random | 261 | 2 | 4.26% | 0.77% | 2.84% |
| BtkV-Cas9 | 4 | 47 | Random | 189 | 1 | 2.13% | 0.53% | 1.96% |
| BtkV-Cas9 | 4 | 30 | Random | 314 | 1 | 3.33% | 0.32% | 1.18% |
| BtkV-Cas9 | 4 | 28 | Random | 180 | 1 | 3.57% | 0.56% | 2.06% |
| BtkV-Cas9 | 8 | 31 | Random | 378 | 0 | 0 | 0 | 0 |
| BtkV-Cas9 | 8 | 29 | Random | 144 | 0 | 0 | 0 | 0 |
| BtkV-Cas9 | 16 | 34 | Random | 121 | 0 | 0 | 0 | 0 |
| BtkV-Cas9 | 16 | 22 | Random | 284 | 0 | 0 | 0 | 0 |
| BtkV-Cas9 | 16 | 29 | Random | 73 | 0 | 0 | 0 | 0 |
| BtkV-Cas9 | 0 | 14 | <24 hpe | 226 | 0 | 0 | 0 | 0 |
| BtkV-Cas9 | 0 | 21 | <24 hpe | 118 | 15 | 71.43% | 12.71% | 20.72%* |
| BtkV-Cas9 | 4 | 15 | <24 hpe | 96 | 0 | 0 | 0 | 0 |
| BtkV-Cas9 | 4 | 26 | <24 hpe | 243 | 2 | 7.69% | 0.82% | 1.03%* |

\*Sex ratio corrected based on ratio of emerged males and females in the injected group.

### Knockout of the B. tabaci white gene by ReMOT Control

For proof of principle gene editing experiments, we chose the *white* gene, which is an ABC transporter protein responsible for transport of ommochrome pigments into the eyes. Null mutants for this gene have white eyes in multiple taxa (20 - 22). Cas9 proteins were complexed with 5 sgRNAs targeting exons 2, 3 and 5 of the gene (Supplementary Table 1). We used saponin as an endosome escape reagent as it was functional in *Aedes aegypti* and *Anopheles gambiae* ReMOT Control editing experiments (9, 10).

Cas9 RNP injections with no targeting peptide or with P2C as the targeting peptide, with or without saponin, did not result in any visibly edited offspring (Table 1). For injections using BtKV as the targeting peptide, white-eye nymphal offspring were recovered from 7 of 14 independent replicate injection experiments (Fig. 2, Table 1). The data showed that mutants were only recovered in treatments without saponin, or the lowest concentration of saponin; higher concentrations of saponin were inhibitory (Table 1). Mutants were recovered in 7 out of 9 independent injection experiments using less than or equal to 4 μg/mL saponin. Finally, females less than 24 hpe with or without saponin generated mutants with higher efficiency compared to females of unknown ages (Table 1). Early nymphal instar mutants had white (colorless) eyes but during development the color changed to a bright orange-red color (Fig. 2) compared to the brown color of the wild-type.

Due to the haplodiploid nature of *B. tabaci*, we are much more likely to observe mutant males as they are haploid. While we can not determine the sex of mutants that did not survive to adulthood, all mutants that did make it to the adult stage were male. We assume that mutants identified in the immature stage were male as well. This can bias actual estimates of editing efficiency if the sex ratio is not 50:50. *B. tabaci* females can actively control the sex ratio of their offspring by controlling which eggs are fertilized, and published estimates of the sex ratio range from 25% to 75% female (23). We examined the sex ratio in our colony 9 replicate times by random sampling and found it to be highly female biased, with estimates ranging from 65% to 85% female, and a average value of 73% female (Supplementary Table 2). When adjusted for sex ratio bias, we calculated an editing rate in male offspring within a range of 0.39%-20.72% (Table 1), depending on whether young females were injected. When females <24 hpe were injected, editing rates in the offspring were >20% in some replicates, high enough for lineage screening by PCR to identify mutant individuals in the case of a gene without an obvious morphological phenotype. Across all replicates using 4 μg/mL saponin or less, injection of approximately 12 females was enough to obtain at least one edited offspring, an editing efficiency almost two-fold greater than observed in mosquitoes (9, 10).

### Analysis of mutants

From the 23 mutant offspring visually identified, we randomly selected 8 for molecular characterization and were able to obtain DNA of sufficient quality for sequencing from 5 offspring (3 white-eye 1st or 2nd nymphal instars, and 2 orange eye male adults. PCR of house-keeping genes was unsuccessful for the other 3 individuals, indicating degraded DNA. We validated gene knockout by PCR and sequencing of the product of the targeted gene locus in these 5 individuals. The sequencing produced multiple peaks for one individual, and so the PCR product was cloned into pJET1.2 and 9 clones were sequenced. This individual (although phenotypically white eye) appeared to be a genetic mosaic, where we detected wild-type alleles and 2 insertion alleles (a 3 nucleotide insertion and a 2 nucleotide substitution). Another white-eye juvenile had a 21bp in-frame insertion (insertion mutant [21bp]). Homology modeling of this mutation predicted that it caused a loop to be cast across the pore region of the transporter, likely blocking the transport function of the gene by steric interference (Fig. 3C). The final white-eye juvenile had a 4bp deletion in exon 3 causing a frame shift of the *white* gene. Similarly, the two orange-red eye males had the same 4bp deletion.

### Heritability of generated mutations

To demonstrate that the mutations generated by ReMOT Control were were heritable in *B. tabaci*, we performed a cross (Fig. 4) between male and female offspring of injected females. The males used in the cross were visibly mutant (white-eye as juvenile, orange-red eye as adults). The females were phenotypically wild-type. All 90 G1 offspring had wild-type eye color which indicates that the parents consisted of a mutant male and wild-type females. Subsequently, G1 females were backcrossed with the same G0 mutant male. The expected ratio of mutant to wild-type offspring resulting from a cross between a haploid mutant male to a hemizygous mutant female is 50:50. We observed 22 mutants and 81 wild-type offspring suggesting a deviation from the expected ratio (Chi-square = 17.84, *P* < 0.0001), Nevertheless, the data demonstrate that the mutation was heritable and thus the germline was edited.

## Discussion

In this report, we demonstrated heritable CRISPR/Cas9 gene editing in *B. tabaci* using ReMOT Control by changing the ovary targeting ligand from P2C to BtKV. P2C was derived from the yolk protein of *Drosophila melanogaster* and was used for gene editing in *Aedes aegypti* (10) and *Anopheles stephensi* (9). In our experiments, P2C did not generate knockouts in *B. tabaci*; thus, we identified a new 13 aa ligand from the native vitellogenin protein of *B. tabaci* to target the RNP for ReMOT Control. This resulted in high efficiency gene editing. In mosquitoes use of an endosomal escape reagent is critical to the success of ReMOT Control (9, 10), and saponin was shown to be highly effective (9). However, in *B. tabaci* we found a chemical endosomal escape reagent was unnecessary, and in fact saponin was detrimental to the process. This result demonstrates that ReMOT Control must be independently optimized for each new species, as parameters that are successful in one system may not work in others.

ReMOT Control transduces Cas9 RNP into the ovaries of females, so it cannot edit the paternal derived chromosome until after the egg has been fertilized. Thus, the maternal chromosome is edited more efficiently than the paternal chromosome (9, 10). *B. tabaci* is haplodiploid, where females are diploid and haploid males result from unfertilized eggs. Because ReMOT control preferentially edits the maternal chromosome, obtained white-eye nymphal instars were likely haploid males that developed from an unfertilized egg with a single edited chromosome. Haplodiploidy is an advantage for ReMOT Control because mutations in the female germline can be immediately recognized in the haploid male offspring. Consistent with this, all G0 mutant adults with orange-red eyes were males. To effectively exploit the haplodiploidy feature, we obtained females less than 24 hpe to inject to limit pre-injection egg development, as well as potentially limiting the chances of mating thus favoring male bias offspring, making it easier to screen for the visually noticeable phenotype.

Our cross experiment demonstrated that mutations generated by ReMOT Control are in the germline and can be passed down to offspring by heredity, rather than just editing the somatic tissues. Mutations were inherited in in deviation of expected Mendelian ratios, possibly due to differential hatching of mutant vs. wild-type eggs.

We observed white-eye nymphs from 1^st^ to early 4^th^ instar and an unexpected orange-red eye phenotype in the mutant late 4^th^ instar nymphs and adults. During the late 4^th^ instar nymphal stage, the eyes of the pharate adult become diffused, heavily pigmented, and distinct (24), which means that the noticeable change in eye color during the late stages of the 4^th^ nymphal instar are the adult eyes. The change in pigmentation in *B. tabaci* is different from the brown leafhopper *Nilaparvata lugens*, another hemimetabolous insect. Mutation of the *white* gene in *N. lugens* yieled a white ocelli and a light red pigmented eye where the eye pigments were consistent throughout the life stages (14). Both ommochrome (brown/black) and pteridine (red/orange/yellow) pigments contribute to eye color in insects (25). We speculate that the *B. tabaci white* gene is responsible for transportation of the brown ommochrome pigments into the eye; with it mutated the red pteridine pigments become visible, explaining the shift in eye color of mutant insects, although further research is required to confirm or refute this hypothesis.

## Conclusion

We have shown in this work that ReMOT Control allows easy and efficient CRISPR editing in *B. tabaci* without the need to inject embryos. The eggs of *B. tabaci* have a pedicel that is embedded into leaf tissue and acts a conduit for water and solute absorption into the eggs for successful embryonic development and hatching (26). This character, along with the exceeding small size of the embryos (∼0.2mm x 0.01mm) and high mortality of injected eggs presents significant challenges for the success of embryo injections. ReMOT Control removes these constrains and significantly expands the ability for any laboratory to apply CRISPR techniques to whitefly research, which will greatly accelerate molecular biology research on this organism and lead to the development of novel control strategies for this economically devastating pest insect. Furthermore, this technology can be applied readily to other whitefly species and to related insect groups, e.g. psyllids, mealybugs, which also include major agricultural pests.

## Acknowledgements

We thank Dr. Angela Douglas for helpful advice regarding whitefly rearing and experiments, Abdulla M.A.J. Alghawas for assistance with laboratory procedures, and Dr. Blake Bextine for facilitating our initial investigations in this area of research. This work was funded by NSF/BIO grant 1645331, NIH/NIAID grant R21AI111175, USDA/NIFA grant 2014-10320, USDA Hatch funds (Accession #1010032; Project #PEN04608), and a grant with the Pennsylvania Department of Health using Tobacco Settlement Funds to JLR.

## Figure legends

**Supplementary Figure 1.**
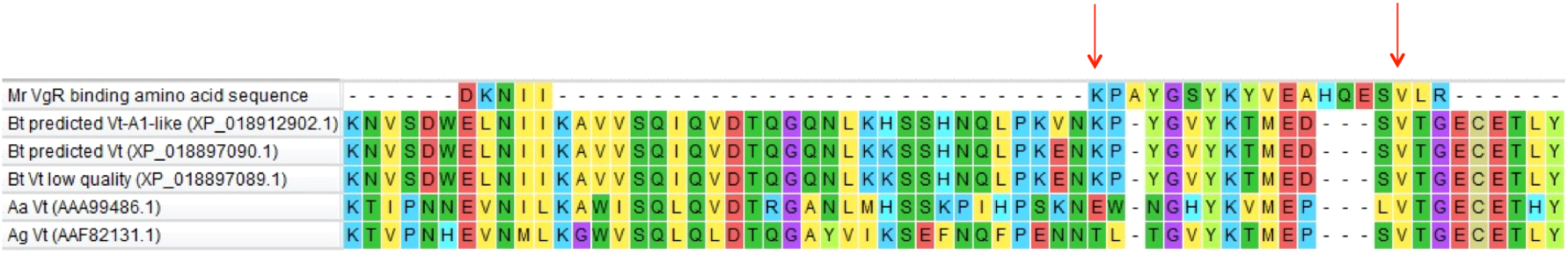
Identification of the BtKV ligand. *Microbrachiumrosenbergii* 24 amino acid sequence identified in Roth et al. 2013, *B. tabaci* predicted vitellogenin-A1-like (XP_018912902.1), *B. tabaci* predicted vitellogenin (XP_018897090.1), *B. tabaci* predicted low quality vitellogenin (XP_018897089.1), *Aedes aegypti* vitellogenin (AAA99486.1), and *Anopheles gambiae* vitellogenin (AAF82131.1) were aligned using ClustalW in MEGA 7.0.26, then manually aligned the “KP” amino acids to match the “KP” of the vitellogenins of *B. tabaci*. As the targeting ligand we chose the sequence from the *B. tabaci* “KP” to the conserved valine (V) (red arrows).

**Supplementary Table 1.**
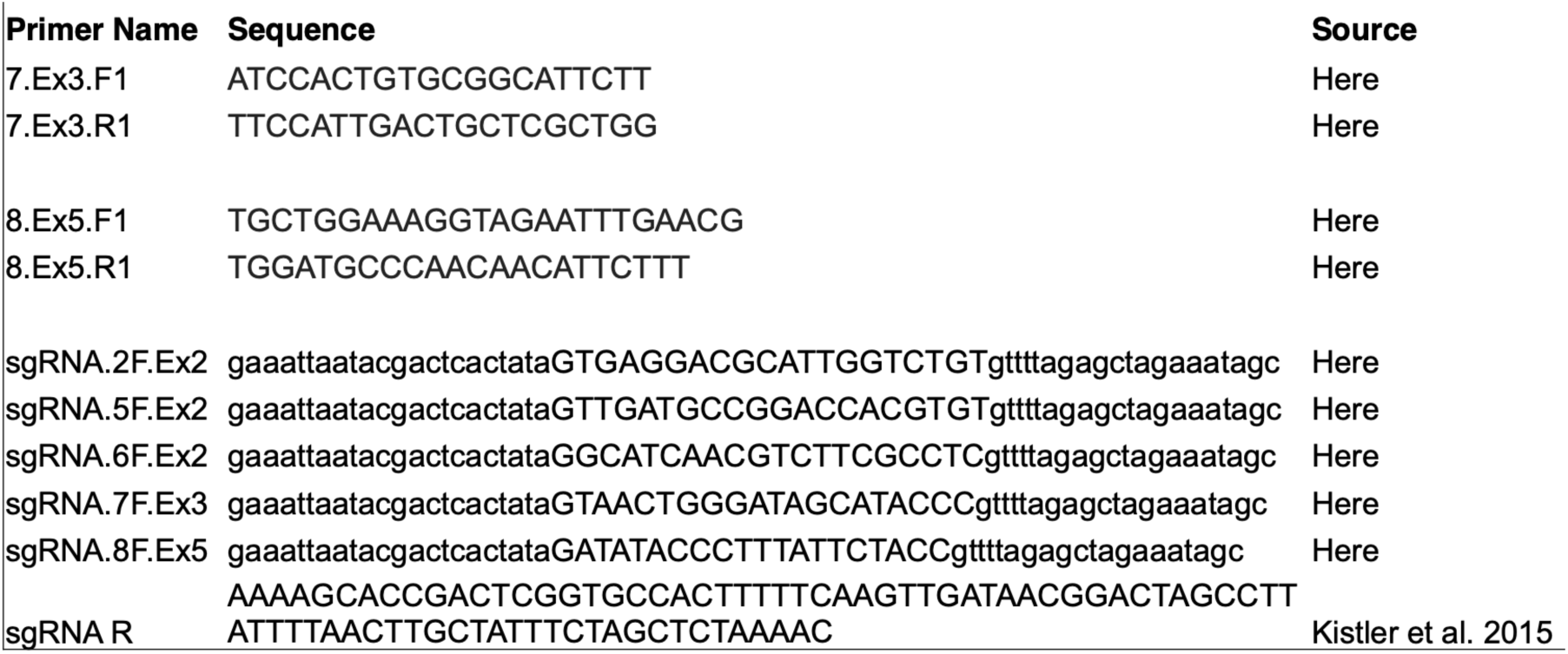
Primer and guide RNA sequences used in this study.

**Supplementary Table 2.** *B. tabaci* colony sex ratio.

| sample | female | male |
| --- | --- | --- |
| 1 | 44 | 17 |
| 2 | 20 | 11 |
| 3 | 20 | 9 |
| 4 | 21 | 5 |
| 5 | 11 | 4 |
| 6 | 22 | 8 |
| 7 | 22 | 6 |
| 8 | 22 | 7 |
| 9 | 22 | 9 |
| Average | 22.7 | 8.4 |
| **Ratio** | 2.7 | 1 |

